# In Poetry, if Meter has to Help Memory, it Takes its Time

**DOI:** 10.1101/2021.03.14.435310

**Authors:** Sara Andreetta, Oleksandra Soldatkina, Vezha Boboeva, Alessandro Treves

## Abstract

To test the idea that poetic meter emerged as a cognitive schema to aid verbal memory, we have focused on classical Italian poetry and on its three basic components of meter: rhyme, accent and verse length. Meaningless poems were generated by introducing prosody-invariant non-words into passages from Dante’s *Divina Commedia* and Ariosto’s *Orlando Furioso*, which were then further manipulated by selectively ablating rhymes, modifying accent patterns or altering the number of syllables. The resulting four versions of each non-poem were presented in a fully balanced design to cohorts of high school educated Italian native speakers, who were then asked to retrieve 3 target non-words. Surprisingly, we found that the integrity of Dante’s meter has no significant effect on memory performance. With passages derived from Ariosto, instead, removing each component downgrades memory by an amount proportional to its contribution to perceived metric plausibility, with rhymes having the strongest effects, followed by accents and then by verse length. Counterintuitively, the fully metric versions required longer reaction times, implying that activating metric schemata involves a cognitive cost. Within schema theories, this finding provides evidence for high-level interactions between procedural and episodic memory.

## Introduction

Poems, nursery rhymes, traditional songs: they are found in every culture and they have been around for ages, well before the advent of writing systems. Sometimes, they have or had the crucial mission of carrying an important message for the listeners: a list to know by heart, an event happening every year, a warning of a potential danger. What do these texts have in common? At least one aspect: they adopt a variety of devices that help hold verbal material in memory.

Human memory can, in fact, fail spectacularly at times. Writing systems have helped safely store verbal information, in a format relatively difficult to tamper with; before, when our ancestors had to rely on their fallible memory, a number of linguistic devices crystallized to help them remember words and verbal material. Cultural transmission, then, has depended for ages on these devices, which in poetry we can broadly refer to as “meter”. These devices may range from the use of repeated metaphors: “rosy-fingered dawn” in Homer [1], to the *ring composition* in the Zoroastrian Yasna [2] to semantic repetition as in Biblical poetry: “In the way of righteousness is life; and in the pathway thereof there is no death.” in Prov. 12:28 (King James Version).

In several Western literary traditions, including the Italian one, the local structure of poetry revolves around the verse, and includes a constant number of syllables, a limited variability in the pattern of accents, a specific organization of rhymes. These components of meter have gradually lost their centrality or at least their perceived necessity over the course of several centuries, but they were in full sway at least between the XIII and the XVI century, from the emergence of modern Italian (so called “volgare”, the language of the people) as an acceptable literary language to the diffusion of the printing press. The *Divina Commedia* by Dante Alighieri and the *Orlando Furioso* by Ludovico Ariosto are two lengthy masterpieces towards the beginning and, respectively, the end of this golden age. With 14.233 verses in the *Commedia* and 38.736 in the *Orlando Furioso*, neither of which contains material which is absolutely necessary to remember in order to carry on with one’s life, it may be asked whether their metric structure still retained a primary memory function, or is already a purely esthetic ornament for cultured readers [3].

Can the role of metrical features be explained from a neurocognitive point of view, with respect to memory? Recently, in the memory literature the notion of schemata, long seen as important (see e.g. [4]) has been discussed again [5] stimulated also by the analysis of its neurobiological basis in rodents [6]. A schema, whether directly functional like those involved in preparing coffee [7] or social/ornamental like rituals of salutations [8], can be considered as a set of regularities that help organize and retrieve information [9]. In this context, we consider the components of meter as schemata which, by encouraging regularities, facilitate the recall of verses. They operate as schemata insofar as they help us recruit, and possibly produce, the next element of a sequence stored in our memory.

In facilitating verbal sequence replay, metrical features appear to be effective with extended “trajectories”, lasting even several verses. These are extended relative to the short trajectories thought to be produced by the phonological loop of Baddeley’s model of working memory, which are presumed to last only a couple of seconds, precisely because of the lack of specific devices that extend their range [10]. To the best of our knowledge, though, the effectiveness of these features has never been quantified. In this study, we aim at measuring the strength of some metric devices. Specifically, we focus on the three main characteristics of classical Italian meter: rhyme, pattern of accents, and verse length.

## Results

First, we established that we could selectively manipulate specific aspects of meter to moderately decrease the “metric plausibility” of meaningless 8- or 9-verse passages derived from classical Italian poetry. Eight Original Non-Poems (ONPs) were generated from as many passages taken from the *Divina Commedia* by Dante Alighieri (four) and the *Orlando Furioso* by Ludovico Ariosto (also four, see Material and Methods), by changing key content words into non-words with the same prosody, thus removing meaning without altering metric structure. Then, for each ONP we generated, with further limited modifications, a Non-Poem without Rhymes (NPR), one with an altered pattern of Accents (NPA), and one with an irregular number of Syllables in 4-5 of its verses (NPS). All 8 original passages were proper (Italian) hendecasyllables with an accented 10^th^ syllable, therefore when adding or subtracting 1 or 2 syllables in the NPS versions, also the pattern of accents was perforce altered, but we attempted to make the alteration less noticeable than the number change, in contrast to the NPA condition in which there were strictly 11 syllables/verse, but the accents followed more unusual patterns. An example of an NPS we used, from the *Commedia*, is presented in Fig.1, and all non-poems are to be made available.

**Fig. 1.**
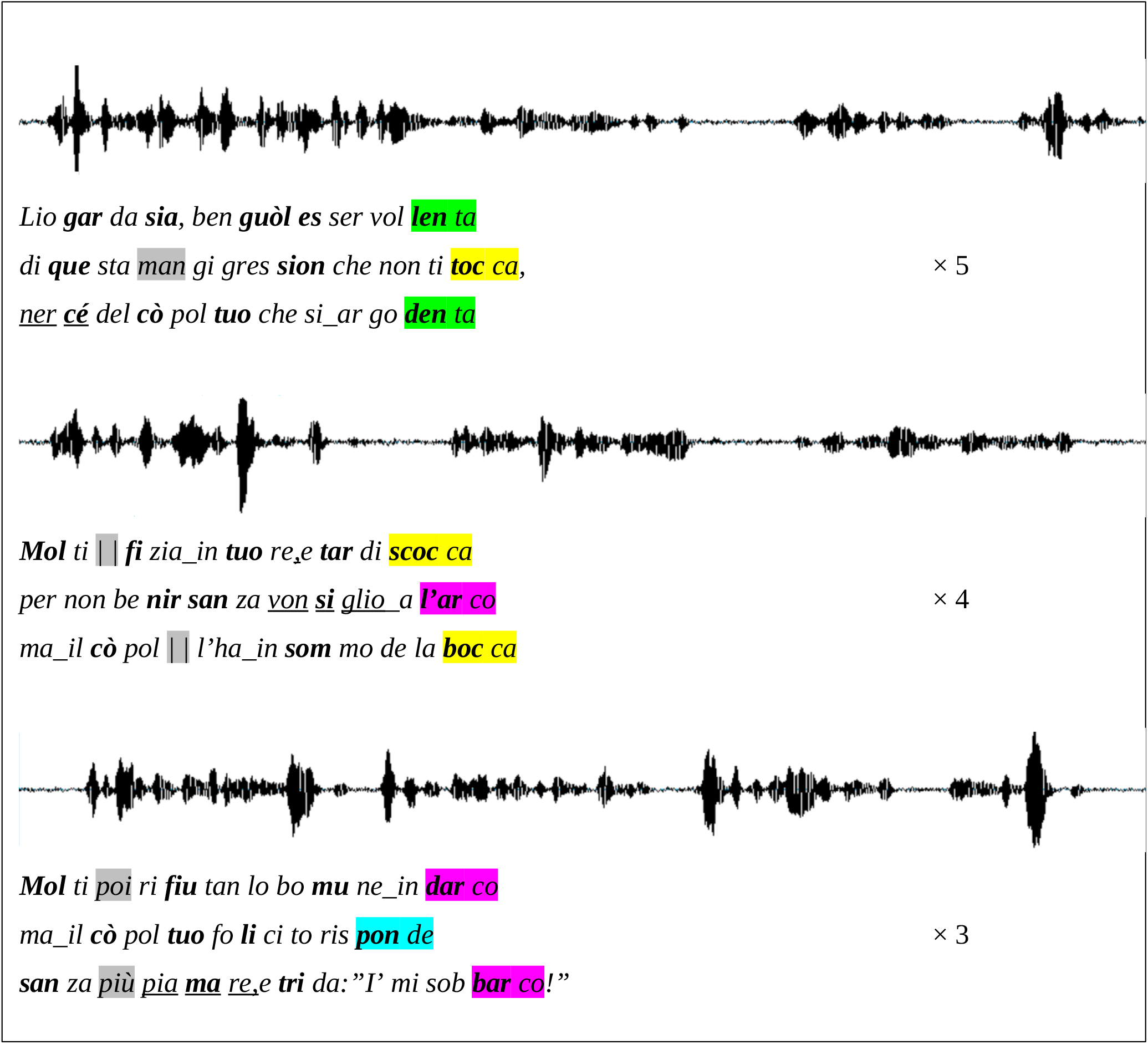
NPS example derived from Purgatorio, canto VI. For each *terzina* (vv.127-135) the NPS text, shown below the sound wave by the professional actor, maintains rhymes, in color, and accents, in boldface, as in the ONP version; whereas overall 3 syllables have been added and 2 taken away, in gray. The underlined non-words were the targets of the memory test; underlined blanks denote *synizesis* (when two syllables are pronounced as one).

Two separate cohorts or rankers, for Dante- and Ariosto-derived non-poems, were presented with a combination of the 4 passages from the same author, one in each of the ONP, NPS, NPS and NPR versions, and were asked to rank them in order of metric plausibility. The fully balanced design (details below) allowed us to extract a passage-independent plausibility score.

### The contribution of distinct components to metric plausibility

Both when derived from passages by Dante and Ariosto, non-sense poems were found most plausible in their fully metric ONP versions, somewhat less when the number of syllables was manipulated (NPS), even less when the pattern of accents was altered in the NPA renditions, and the least when rhymes were removed, NPR. Remarkably, however, differences in the plausibility index are shown in Fig.2 to be quite limited, making the fully balanced design essential. The variance was particularly limited in passages from the *Commedia*, which may be due to Dante’s taking more liberties with the meter he had adopted (the same hendecasyllables as Ariosto, but in *terzine* rather than *ottave*). To quantify this perception, at least in relation to accent patterns, which are more accessible to analysis, we applied two independent measures of accent variability to the 4 original passages by each author.

**Fig. 2.**
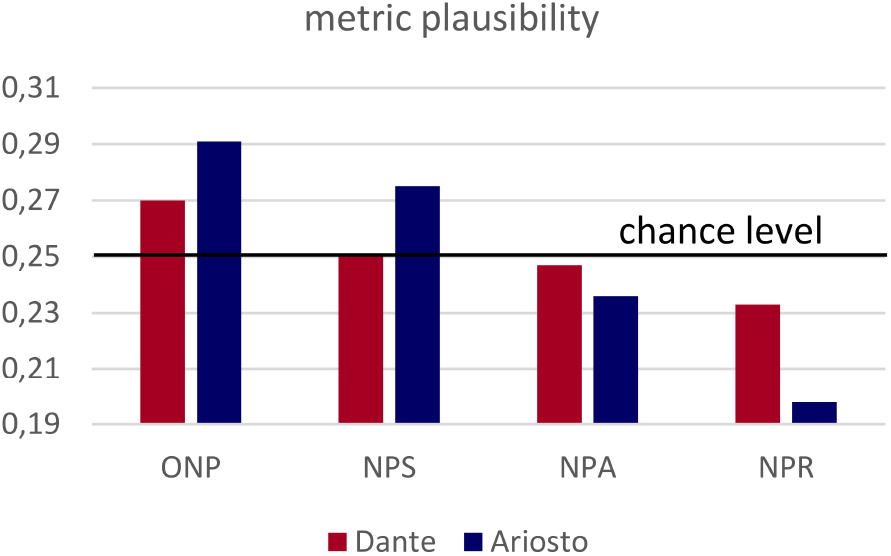
Relative metric plausibility. The different versions of the same 4 passages from the *Divina Commedia* (red) and *Orlando Furioso* (blue) were ranked in the same order, but the plausibility index (see Methods) is more spread out for Ariosto.

Dante appears to be slightly more variable in his accent patterns relative to Ariosto, but the main observation that can be gleaned off Fig.3 is that both poets are far from using a fixed pattern, utilizing over half of the maximum entropy they had available in terms of accenting those passages.

**Fig. 3.**
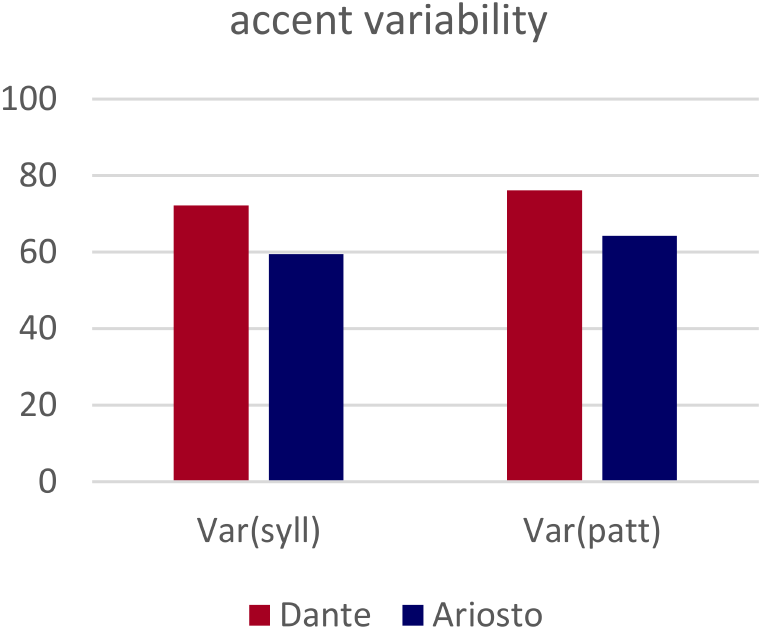
Variability in the pattern of accented syllables in the 8 original passages by the two authors. Two independent entropy measures of variability, per syllable and per verse (see Methods) are both normalized to range from 0 (a single fixed pattern) to 100% (i.e., each syllable in the verse is accented half the time; or each verse follows a different accent pattern).

### Meter can facilitate memory

Does such loose structure help remember individual words? Table 1 and Fig.4 (upper) show that it does, only for the non-poems derived from Ariosto’s octaves. Twenty-four subjects per author were asked, one day after repeatedly listening to one version of each passage, to remember non-word targets out of 3 alternatives, upon listening to the non-poem with selected non-words muted. There were 3 such targets in each non-poem. While in the case of those derived from passages in the *Divina Commedia* the overall number of correct responses per condition was unrelated to its metric plausibility (r^2^=0.04), seemingly fluctuating as much as the correct responses to the first, second and third query taken alone (Table 2), for the passages from *Orlando Furioso* the correlation with metric plausibility was remarkable (r^2^=0.98) and highly significant (p<0.01).

**Table 1.** Correct responses out of a total of 24 participants, for the first, second and third query.

|  | Dante |  |  |  | Ariosto |  |  |  |
| --- | --- | --- | --- | --- | --- | --- | --- | --- |
|  | NPR | NPA | NPS | ONP | NPR | NPA | NPS | ONP |
| 1 | 5 | 15 | 13 | 10 | 11 | 16 | 19 | 13 |
| 2 | 13 | 13 | 9 | 10 | 9 | 10 | 9 | 15 |
| 3 | 15 | 17 | 16 | 12 | 11 | 8 | 12 | 14 |
| All | 33 | 45 | 38 | 32 | 31 | 34 | 40 | 42 |

**Fig. 4.**
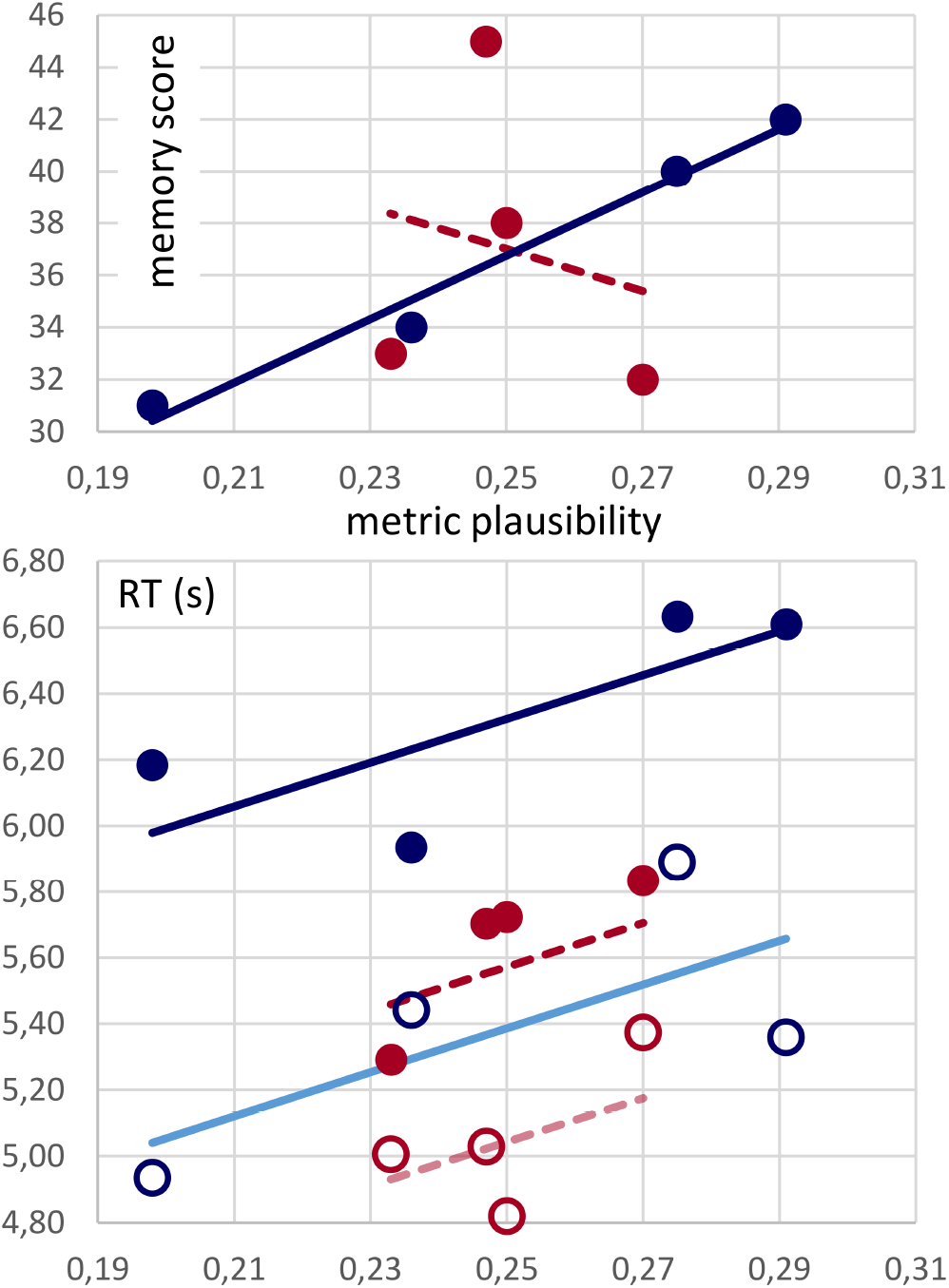
Memory and reaction times both increase with metric plausibility. (Upper) Overall correct responses (out of 72) for each condition, ordered in terms of their metric plausibility, as in Fig.2, for passages from Dante (red) and Ariosto (blue). (Lower) Reaction times (in seconds) for correct (circles) and wrong responses (dots) are regressed against plausibility for each author, with a single slope parameter. The slope is significant and similar to that characterizing the Ariosto data alone, whereas it is denoted with a dashed line for the Dante data, because the latter would not produce a significant correlation on its own.

Interestingly, the total score of the two cohorts was nearly identical, 147 for Ariosto and 148 for Dante, out of a total of 288 (24×4×3).

### Meter helps, but not for free

The analysis of reaction times helps interpret the above results. As shown in Fig. 4 (lower), overall it took longer for participants to pick a wrong answer over the correct answer (on average, 733ms more), and it took longer for participants tested with Ariosto, relative to those tested with Dante, to respond (on average, 547ms more). Most importantly, in each of the 4 types of trials above, the more metrically plausible the passage, the longer the reaction time. However, the trend is significant only with Ariosto, if data from the two authors are analyzed separately, and it is significant overall (p<0.004) with a slope mainly determined by the Ariosto data, if analyzed together, as shown in Fig.4. The slope for the Dante data alone would be higher, but not significant, likely because of the limited plausibility range spanned.

These findings suggest that processing meter in order to help retrieve a non-word heard the day before has a cognitive cost, and takes of the order of hundreds of ms extra time, depending on exactly how much meter there is “used up” in the process. For passages derived from Dante, it appears that although outwardly the metric structure is essentially the same (with the slight qualification reported in Fig.2, and the note that a passage is a sequence of 3 *terzine* rather than a single *ottava*), meter is used less, and the very same memory performance is attained on average in less time.

## Discussion

The connection between meter and memory is not new to cognitive science: in a seminal book Rubin described oral traditions and the linguistic devices they use, highlighting in particular their role in memory as limiting the choice (one could say the entropy, [11]) of larger units: by indicating a specific word ending, for instance, choices will be limited to those words which have the same ending, if a rhyme is expected [3].

In music, Schulkind has investigated memory mechanisms by having participants listen to well-known and novel songs which were altered in their rhythm. Results showed that unaltered versions were identified significantly better than the altered ones, and this applied to both known and novel songs [12].

Analogously, Sachs investigated the retention of semantic and syntactic information in discourse by having participants listen to short prose stories. By selectively manipulating the meaning or the syntactic form of a target sentence, she could show that meaning is remembered, in prose, better and for longer that meaning-irrelevant sentence form [13].

In a similar design, Tillman and colleagues have tested short term memory in prose and poetry. Also in this case, a sentence, considered the target, was changed in its form or in meaning. While with prose memory for surface characteristics declined over time, as expected, the same did not happen with poetry, for which form, in addition to content, was efficiently retrieved [14].

With this study, we had hoped to be able to quantify, in rather absolute terms, the contribution of different aspects of meter to memory retention, using “material” from the classical period of Italian literature, before the advent of the printing press diminished the perceived value of memorability *per se*, and promoted the further ritualization of the written verse into a primarily esthetic construct. The results belied our naïve expectation, in that meter seems to be “perceived” much more (in terms of our plausibility index) in passages derived from Ariosto than in those from Dante, and to contribute to memory in the former but not in the latter. Yet the meter employed by the two authors is nearly identical, with a discrete difference in the concatenation of hendecasyllables (in *terzine* in the *Divina Commedia*, in *ottave* in the *Orlando Furioso*) and a presumably small quantitative difference in the variability with which the common meter is used (Fig.3). Therefore, one would expect that the listener, or the reader, activates the very same cognitive schema, at least locally, within the few verses of a single *ottava* or 3 *terzine*.

The time the subjects from the two statistically indistinguishable cohorts needed to react to the memory tests suggest an account of the main finding: the metric schema is the same, but it is activated to a different extent. Somewhat counterintuitively, it appears to be activated less with Dante, an author with whom most people who have been in high school in Italy are rather familiar, than with Ariosto, who has been relegated, especially in the last few decades, to a marginal niche in standard Italian curricula. This appears to discount a possible interpretation of this difference, i.e., that we are seeing two competing effects, whereby both congruence and incongruence with established schemata can enhance memory, the latter a novelty effect [15]: novelty presupposes the activation of the schema it contradicts. An alternative interpretation is that Dante’s verses are just more interesting, and tend to focus one’s attention to other aspects than the components of meter. Even if this interpretation were to be shown to be correct, it is quite surprising that it would apply, in our paradigm, to verses that have been deprived of their meaning. Moreover, our replacing several of the key words chosen by Dante with our untalented choice of non-words would have been expected to remove other potentially useful devices from the poet’s bag-of-tricks, like alliteration, onomatopoeia, use of liquids, of newly crafted words, etc. [16].

Can the hypothesis of differential schema activation be tested experimentally? In principle, yes, and one approach would be by looking at evoked response potentials (ERPs), which have been widely used to reveal brain signals that reflect violations of expectation, whether (in the language domain) syntactic, semantic, or just phonological [17][18]. With poetry, there have been e.g. ERP studies of aesthetic appreciation and ease of processing [19] and of brain activity during poem composition [20]. For an experimental design like ours, however, one challenge is how to obtain the large number of trials per condition needed in order to obtain valid ERP measures. Another one is to what extent one can rely on single ERPs to characterize a heterogeneous variety of metric components. It is possible that addressing both challenges will require a change in perspective from whole brain dynamics to one which articulates the cortex into a plurality of interacting local networks, as embodied e.g. by the Potts model [21]. Distinct processes, among the many that concur to the overall perception, appreciation and memory of a poem, including the components of meter, are likely reflected differentially in the dynamics of distinct cortical networks, just like other, better studied types of memory such as episodic and spatial memories [22], which have stimulated theories about the interactions between the medial temporal lobe and medial prefrontal cortex [9]. Meter, with its multiple components, indicates the need to go beyond the somewhat coarse distinctions available to neuropsychology, what is at the moment accessible only through mathematical models [23], with which one can study forms of partial coherence among multiple local networks, reminiscent of that of systems unable to attain long-range order [24].

While partial coherence might seem to detract from the wholeness attributed to conscious processing [25], it is entirely consistent with the idea of a mixture of automatic and controlled components concurring to memory encoding and retrieval [26]. With meter, the notion that different schemata might be activated only optionally, at times, and then partially and incoherently with others, and when activated might offer only an incremental contribution, suggests a more nuanced take on high-level cognition in general. Using many filters to interpret reality, and to a variable degree, implies check-and-balances and minimal recourse to prevailing or dominant schemata, those that often reflect biases or prejudice.

## Material and Methods

We extracted passages from two masterpieces of Italian literature: the *Divina Commedia* by Dante Alighieri (1265-1321), and the *Orlando Furioso* by Ludovico Ariosto (1474-1533). From the latter we chose *ottave* (octaves, stanzas of 8 verses) from *canti* XIII, XV, XIX and XXX, and one from *canto* I for training subjects, while from the former we selected sequences of 3 consecutive *terzine* (hence 9 verses) from two *canti* from *Inferno* (XXIV for the experiment, V for training), two from *Purgatorio* (VI and XVI) and one from *Paradiso* (XXVII).

### Poem manipulation

The original texts were manipulated in a number of different ways. First, most content words were converted into non-words in order to eliminate discernible semantic content, hence semantic effects on memory; an effort was made to change phonemes with similar ones, while maintaining the original prosody. Function words were not modified. This applied equally to all texts, and resulted in “original non-poems” (ONPs).

The second stage of manipulations focused on metrical patterns. In particular, we created three conditions:

1. a condition where we eliminated rhymes (“NPR” – Non-Poem without Rhyme)
2. a condition where the accent patterns of 4-5 verses per passage were replaced with less standard ones (“NPA” – Non-Poem with modified Accents). To validate proper original accents we consulted with an expert scholar for *Orlando Furioso*, whereas for *Divina Commedia* we referred to the “Archivio Metrico Italiano”, a database collecting masterpieces of Italian literature with their accents annotated [27]
3. a condition where the number of syllables per verse, which in the ONPs were strictly 11 throughout (regular hendecasyllables) was altered again in 4-5 verses, to 9, 10, 12 or 13 - (“NPS” – Non-Poem with wrong numbers of Syllables)

These manipulations were applied to all texts.

For the experiment, every subject was administered four texts in total, by the same (original) author: one per *canto* and one per condition. Therefore, twenty-four combinations were created.

All texts were then recited by a professional actor and audio recorded.

An example of an NPS we used, from the *Commedia*, is presented in Fig. 1 together with its original spectrogram, and all non-poems will be made available separately.

### Ranking

This experimental material was used to conduct an online survey about how these manipulated poems were perceived by a group of Italian native speakers. Participants were asked to listen to the four conditions and give a ranking of preference, from the one that sounded the most “poetically plausible” to them, to the one they perceived as the strangest.

#### Subjects

62 people participated in the online survey for Ariosto (F=32, M=30, mean age = 29.06, sd= 8.13) and 65 people for Dante (F=35, M=30, mean age = 26.48, sd = 6.26). Part of either cohort were the participants in the main experiment below, but tested with the other author, and they were asked to complete this survey after the end of the second session of the main experiment. Another group of participants was recruited through the online platform Prolific [28]. This last group was compensated with 5 euros. We had aimed for 72 rankers in each cohort, but had to exclude some *a posteriori*, who failed to complete the ranking in full.

#### Experimental design

The online survey was designed with the open source toolkit Psytoolkit [29]. After an example, presented as training also in the main experiment below, they listened to the four poems one at a time. Every poem was associated with a name, in order to help participants refer to that specific condition. If they wanted, they were allowed to listen again and again to the same poem before proceeding to the next.

At the end they were asked to rank the four poems: from the one they perceived as the best, to the one that sounded worse to them. From the rankings by all participating subjects we extracted an average index of metric plausibility by assigning a value 0.6 to the first –ranked condition (e.g., NPA), 0.3, to the second, 0.1 to the third and 0 to the fourth. The logic behind this assignment is that subjects occasionally reported being unsure as to which passage sounded the strangest. The rankings were collapsed across *canti*, with the relatively large number of participants ensuring approximately even sampling (each *canto* was presented originally 18 times per condition, which came down to 16+/-2 after the exclusions). As a result, the average metric plausibility of each condition could in principle range from 0 to 0.6, but in practice was much more restricted, particularly with passages from Dante, to values around the average of 0.25 (see Fig.2).

## Memory experiment

### Subjects

Forty-eight native Italian speakers who had been exposed to Italian literature through one of the national high school curricula were recruited for the main experiment. Half of them were administered material from Ariosto (F= 15, M=9, mean age= 26.34, sd=4.02), the other half from Dante (F=15, M=9, mean age= 26.12, sd=3.61).

Participants were recruited through the SISSA recruiting platform and social media. None of them had a previous history of psychiatric or neurological illness, learning disabilities, nor hearing or visual loss. They were asked to participate in a study on memory and poetry, which would have involved them for two consecutive days, for about 30’ the first and about 10’ the second. Due to the pandemic situation they were asked to be connected remotely with their own devices. All participants signed the consent form before taking part in the study. Their participation was compensated with 15 euros.

### Experimental design

The experiment was designed with Psychopy Builder [30]. It included a *study phase* of about 30’ the first day, and the *test* of about 10’ the second day.

We aimed at an almost exclusively auditory experiment, in order to assess how memory relies on meter if listening is the only available channel to learn from [31]. Indeed, the material included audio files only, with the sole exception of written material when a *fill in the gap* task appeared.

Besides the four passages, verses from 2 other *canti* were used for training, as indicated above. However, these verses were presented only in the ONP condition, leaving meter intact.

Every passage, including the training, was associated with an image, taken from among Gustave Doré’s illustrations of the *Divina Commedia*. The images were consistent for the same *canto* in different conditions and were intended to help engage memory without, at the same time, biasing the linguistic material (images will be made available separately).

Every poem, including the training, was presented divided in three consecutive portions.

### Study phase

The study phase started with the training ONP. First, participants listened once to all verses. Then they listened to the first part (3 verses) repeated five times, with a 3s pause between repetitions. Afterwards, another repetition of the same verses followed, but this time a non-word was muted. The task for the participant was then to retrieve the correct non-word. Three written alternatives appeared on the screen and the participant had to read aloud his/her choice.

After this training, each of the experimental passages was played, in a separate block, in the one of the 4 versions that had been assigned to that subject in the design. Then, after listening once to the entire poem (9 verses in Dante, 8 verses in Ariosto), participants listened to the first part (3 verses for both authors) 5 times. Next they had to complete the task with the muted non-word.

The same happened with the second part of the poem, repeated 4 times. In this case the audio started from silence with the first part ramping up linearly in intensity, until it continued smoothly into the second part at the standard volume. Therefore, this allowed them to have a feedback about the test they just completed.

For the third part they just listened to the repetitions (3 verses for Dante, 2 verses for Ariosto; × 3), starting now with an acoustically smoothed version of the second part, but there was no test.

The alternatives to the correct non-word were chosen by maintaining the same number of syllables, and the same accent. Typically, stems and intermediate vowels or consonants changed. Target non-words were generally in the third verse, aiming not to overload working memory from the moment they listened to the silent word until the test time. In the cases where this was not possible (because there were no appropriate non-words in the third verse, for instance) a non-word in the second verse was chosen, if possible towards the end. Notably, for every text we chose options which were consistent among all conditions, allowing a fair comparison in the results.

### Test phase

The following day participants were connecting at the same time, so 24 hours had passed. They listened directly to the test parts of the text, so the verses with a muted non-word, and they were asked to identify the muted target in all three parts, including then the third one. For the first and second part, tested also the previous day, target non-words were different. After completing the three tests per text, they listened to the entire poem, to receive feedback.

## Supporting information

online Appendix with Suppl Figs.1-5

## Acknowledgements

We are grateful to the HFSP collaboration RGP0057/2016 on Analog computations underlying language mechanisms, in particular to Yair Lakretz, with whom the original idea for this study was discussed, and to Elisa Ciaramelli, Rodolfo Zucco, Sergio Bozzola, Andrea Tabarroni and Sara Alzetta, who offered their different perspectives.

## Contributions

SA and AT conceived the experiment and generated the non-poems. SA and VB implemented it in Psychopy and Psytoolkit, and SA run it, assisted by OS with Prolific and with data analysis. SA and AT have written the report with inputs from VB and OS.

