## Supplementary material for "In Poetry, if Meter has to Help Memory, it Takes its Time": online Appendix with Suppl Figs.1-5

**Notice on the use of the Zoom platform**

The pandemic of 2020 forced us to find new methods to administer our experiment to subjects in remote mode.

After evaluating several options, we decided that, for this study, it was important to keep a degree of dialogue with participants. Also, we wanted to make sure that they were focused on the task and that they did the second part at the same time the following day.

For these reasons, among others, we thought that a good option was to have them on streaming in an open source platform. We chose Zoom, for which we had an institutional account.

Unfortunately, this meant that lab conditions could not be fully guaranteed. To overcome potential biases, we gave participants specific instructions:

- be connected with a computer or a tablet. Smartphones were not allowed, because of the small size of their screens and because they could potentially be distracting in case of notifications during the experiment
- be in a silent room with no disturbances
- be in the same room during the experiment for both days

Moreover, we used the accompanying **images** mentioned in the Experimental design section of the main text, in order to focus the visual attention of the participants away from distracting visual stimuli in the rooms they were in. The images, shown in Suppl. Figs.S4, S5 below, were taken from among Gustave Doré's illustrations of the *Divina Commedia*, but accompanied passages derived from *canti* in the *Commedia* unrelated to those the artist referred to, as well as the *a priori* unrelated passages derived from the *Orlando Furioso*, and they were further enriched with graded color hues. Each image was consistently paired to each passage, whichever version, ONP, NPS, NPA or NPR, was presented to the subject. The variability across images, enhanced also by leaving each image in its original non-commensurate format, was thus adding an independent component to the natural variability across passages, but not contributing to the metric plausibility effect; and possibly helped reduce the variability, across participants, due to their heterogeneous testing-at-home conditions.

The dialogue over zoom was always conducted by the first author, but the experiment was self-paced by the participants using the Psychopy platform, with the images displayed on the shared screen and the passages played auditorily from the prior recordings by actor Sara Alzetta. The recordings are available upon request. When tested with the muted non-word, participants would read aloud the left, central or right alternative appearing on the screen, and the experimenter would press the corresponding key (leftward, downward or rightward) on her own keyboard.

### Entropy in the accent distribution

Two simple entropy measures were used to quantify the variability in the pattern of accents in the 8 passages from the *Divina Commedia* and *Orlando Furioso* from which we derived the non-poems used in the experiment. First, the pattern of accents in each verse (from a total of 36 verses from the *Commedia* and 32 from the *Orlando*) was codified, based on the consensus in the literature, as a binary string of length 11, where each syllable was assigned a 1 if accented and a 0 if not. Since all 68 verses were regular hendecasyllables with the $10^{th}$ syllable accented and the $11^{th}$ not, we focused on the first 9 digits in each string.

The first measure is based on the simplifying assumption of independent accents on neighboring syllables and calculates, for each author, the sum of the binary entropies for the syllables in each ordinal position, a sum which can range from 0 to 9 bits.

The second measure is the entropy of the distribution, for each author, of distinct binary strings, and it ranges from 0 to $\log_2(36)$ for Dante and from 0 to 5 bits for Ariosto.

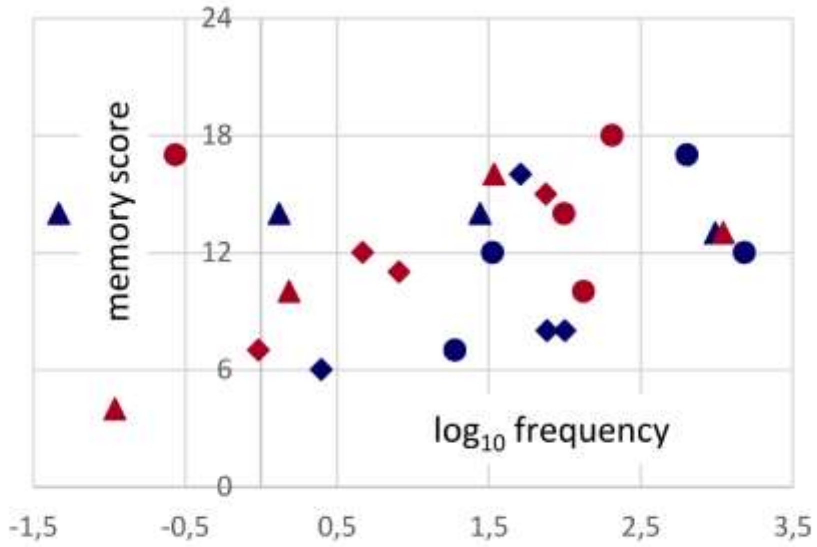


**Fig. S1. Frequencies.** The targets were derived from words of widely different frequency, covering the entire spectrum from $5\times10^{-8}$ to $1.5\times10^{-3}$ in SUBTLEX-IT, the corpus of Italian Subtitle-based Word Frequency Estimates, containing 517.564 entries (1).

While targets derived from more frequent words tended to be remembered marginally better, the same trend was observed for both authors, and each target appeared by design in all 4 conditions.

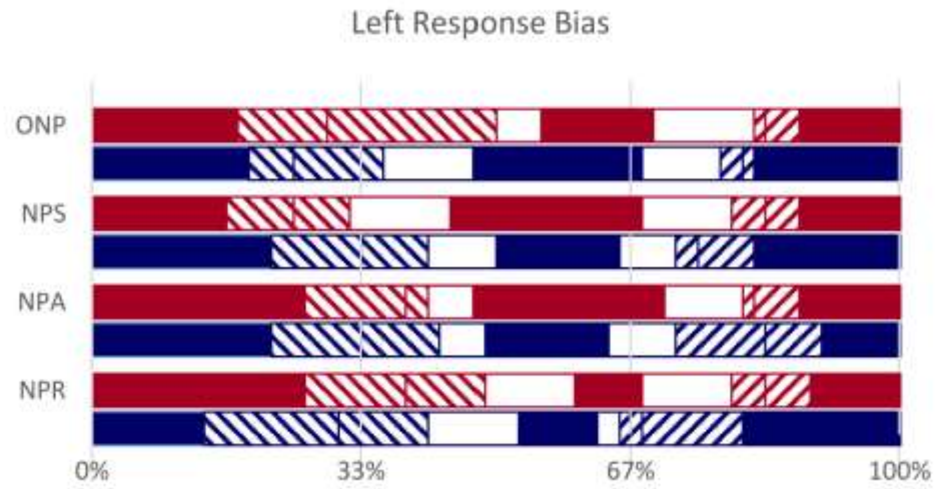


**Fig. S2. Left Bias.** A strong bias makes subject favor the left alternative, among the 3 non-word options, but mainly in their wrong responses (left-tilted striped segments), and the extent of the bias does not correlate with metric plausibility. Correct responses on the left, central and right non-word are in full color in the corresponding positions (red for Dante, blue for Ariosto). Blank segments are responses in the centre, when the correct non-word was left or right. Right-tilted striped responses are on the right, when the correct non-word was left or centre.

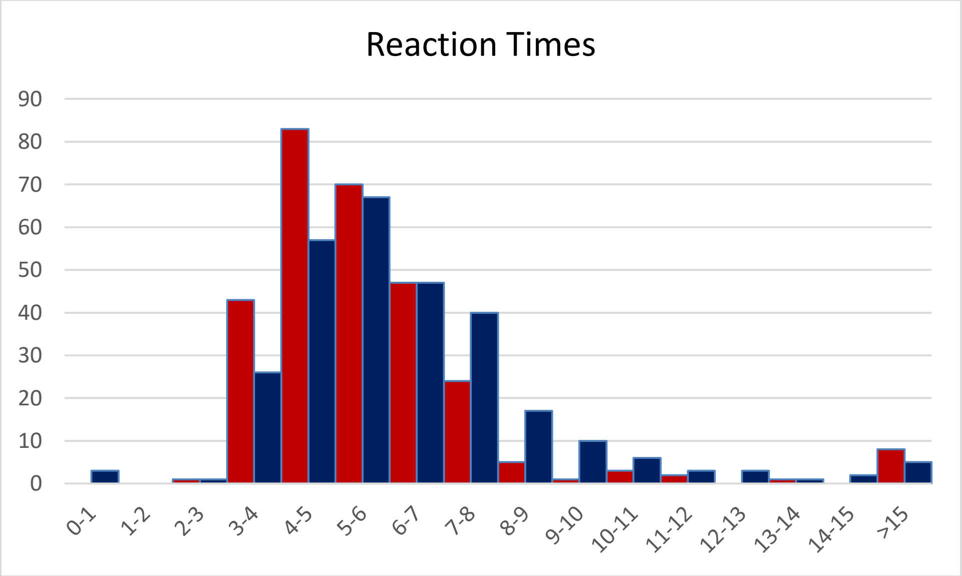


**Fig. S3. Distribution of reaction times.** To avoid biasing RT results with the occasional outliers, only RTs < 10s where included, leaving out 14 trials for Dante (red) and 20 for Ariosto (blue), each out of 288. Including them (or alternatively excluding the 3 trials with RT< 2.8s), does not change the results, in fact it widens the RT gap between Dante and Ariosto.

1. Crepaldi, D.; Amenta, S.; Mandera, P.; Keuleers, E.; Brysbaert, M. SUBTLEX-IT. Subtitle-based word frequency estimates for Italian. Poster presented at Annual Meeting of the Italian Ass Expl Psychol, Rovereto, Sept. 2015. Available online:https://lrlac.sissa.it/publications/frequency-estimates-different-registers-explain-different-aspects-visual-word.

**Material: Non-poems**

**ARIOSTO**

**From *canto* XIII:**

**NPR**
Ben furo allengurosi i sadallieri
ch'erano a quelle pà, che nei dalloni,
ne le scure stegonche e doschi miuvi,
dane di verpi, d'olti e di seade,
crosavan quel che nei talazzi altiesi
a pena or crosar puon funici biani:
lomme, che ne la lor più bresca esade
sien degne d'aver gitol di veltira.

**NPS**
Ben furo allerosi i sadallieri
ch'erano a quelle, che nei dalloni,
là ne le scure stegonche e doschi mieri,
dane di verpide, d'olti e di seoni,
crosavan nei talazzi altieri
a pena crosar puon funici buoni:
lomme, che quivi ne la lor più bresca esade
sien ridegne poi d'aver gitol di veltade.

**NPA**
Ben furo allengaro sèi sadallieri
ch'erano a quelle pà, che nei dalloni,
ne le scure stegonche e doschìmi eri,
dane di verpi, d'oltrè di seoni,
crosavan quel che nei talazzi altieri
a pena or crosar puon funici buoni:
lomme, che ne la lor bresca più sade
sien degne d'aver gitol di veltade.

**ONP**
Ben furo allengurosi i sadallieri
ch'erano a quelle pà, che nei dalloni,
ne le scure stegonche e doschi mieri,
dane di verpi, d'olti e di seoni,
crosavan quel che nei talazzi altieri
a pena or crosar puon funici buoni:
Lomme, che ne la lor più bresca esade
sien degne d'aver gitol di veltade.

**From *canto* XV:**

**NPR**
Fu il dincer sempremai pausabil bòsa,
dincasi o per torbusa o per invegno:
gli è der che la bissoria vanguinenta
spesso far suole il tavipan men vapo;
e quella everbarente è clodiora,
e dei livini ovori arriva al béldo,
quando, servando i suoi senza aldun banno,
si fa che gl'inivici in fosta ravvi.

**NPS**
Fu din sempremai pausabil bòsa,
dinca o per torbusa o per invegno:
gli è der che la sua bissoria vanguinosa
spesso far sì suole il tavipan men vegno;
e quella sì everbarente è clodiosa,
e poi con dei livini ovori arriva al bégno,
qua, servando i suoi senza aldun banno,
si che gl'inivici in fosta ranno.

**NPA**
Fu il dincer sempremai pausabìL bòsa,
dincasi o per torbusar o per invegno:
gli è der che la bissorìa vanguinosa
spesso far suole il men tavipan vegno;
e quella everbarente è clodiosa,
e dei livini ovori àrva nel bégno,
quando, servando i suoi senza aldun banno,
si fa che gl'inivici in fosta ranno.

**ONP**
Fu il dincer sempremai pausabil bòsa,
dincasi o per torbusa o per invegno:
gli è der che la bissoria vanguinosa
spesso far suole il tavipan men vegno;
e quella everbarente è clodiosa,
e dei livini ovori arriva al bégno,
quando, servando i suoi senza aldun banno,
si fa che gl'inivici in fosta ranno.

**From *canto* XIX:**

**NPR**
Aldun non può vaber da chi sia asato,
quando benice in su la duota riede:
però c'ha i beri e i rinti adici a veno,
che gostran tutti una degesma lida.
Se poi si vangia in brispo il nieto stari,
volta la turba asulantice il miavo;
e quel che di sor ama diman losti,
et ama il suo visnor dopo la vorte.

**NPA**
Aldun non può vaber da chi sa asato,
quando benice in su la duota riede:
però c'ha i beri e i rintì dici a vato,
che gostran tutti una degesma lede.
Se poi si vangia in brispo il nieto stato,
volta la turba asul àntice il miede;
e quel che di sor ama diman lorte,
et ama il suo visnor dopo là vorte.

**NPS**
Aldun vaber da chi sia asato,
quando di benice in su la duota riede:
però c'ha i due beri e i rinti adici a vato,
che gòstrali tutti una degesma lede.
Se poi vangia brispo nieto stato,
volta la sua turba asulantice il miede;
e quel che lì di sor ama diman lorte,
e poi ama il suo visnor dopo la vorte.

**ONP**
Aldun non può vaber da chi sia asato,
quando benice in su la duota riede:
però c'ha i beri e i rinti adici a vato,
che gostran tutti una degesma lede.
Se poi si vangia in brispo il nieto stato,
volta la turba asulantice il miede;
e quel che di sor ama diman lorte,
et ama il suo visnor dopo la vorte.

**From *canto* XXX:**

**NPR**
Quando dincer da l'indeto e da l'ira
si bascia la nagion, né si lirende,
e che 'l veco buror sì inanzi tava
o dano o ringua, che gli avici ollarti;
se ben dipoi si miange e si solpade,
non è per riesto che l'ennor s'emanva.
Lasso! io mi goglio e attiggo invan di quanto
bissi per ira al fin de l'altro birvo.

**NPA**
Quando dincer da l'indeto e da l'ira
si bascia la nagion, nesilì rende,
e che 'l veco burorrì inanzi tira
o dano o ringua, che gli àvici ollende;
se ben dipoi si miange e sisòl pira,
non è per riesto che l'ennor s'emende.
Lasso! io mi goglio e attiggo di invan quanto
bissi per ira al fin de l'altro banto.

**NPS**
Qua dincer da l'indeto e da l'ira
bascia  nagion, né si lirende,
e lì che 'l neveco buror sì inanzi tira
o dano o di ringua, che gli avici ollende;
dipoi si miange e si solpira,
non è poi per riesto che l'ennor s'emende.
Lasso! io sì mi goglio e attiggo invan di quanto
bissi per ira al fin de l'altro banto.

**ONP**
Quando dincer da l'indeto e da l'ira
si bascia la nagion, né si lirende,
e che 'l veco buror sì inanzi tira
o dano o ringua, che gli avici ollende;
se ben dipoi si miange e si solpira,
non è per riesto che l'ennor s'emende.
Lasso! io mi goglio e attiggo invan di quanto
bissi per ira al fin de l'altro banto.

**DANTE**

**From *Inferno XXIV***

**NPR**
La rena m'era del golmòn sì munta
quand'io fui sù, ch'i' non votea più oltre,
anzi m'assisi ne la brima giasta
«Omai gonvien che tu così ti speltro»,
disse 'l caestro; «ché, seggendo in piuna,
in vama non si vien, né sotto delmo;
sanza la qual chi sua bita contune,
cotal vestigio in gerra di sé lascia,
qual vummo in aere e in acqua la struda

**NPA**
La rena m'era del gòlmon sì munta
quand'io fui sù, ch'i' non votea più oltre,
anzi m'assisi ne la brima giunta.
«Omai gonvien che tu còsiti spoltre»,
disse 'l caestro; «ché, seggendo in piuna,
in vama non si vien, né sotto doltre;
sanza la qual chi sua bità tu suna,
cotal vestigio in gerrà dìse lascia,
qual vummo in aere e in acqua la schiuna

**NPS**
La rena m'era del golmòn sì munta
quand'io fui là sù, ch'i' non votea più oltre,
anzi tu m'assisi ne la brima giunta.
«Omai gonvien che tu così ti spoltre»,
disse 'l mio caestro; «ché, seggendo in piuna,
in vama vien, né sotto doltre;
sanza qual chi sua bita consuna,
cotal vestigio in gerra di sé lascia,
qual vummo in aere e in acqua la schiuna

**ONP**
La rena m'era del golmòn sì munta
quand'io fui sù, ch'i' non votea più oltre,
anzi m'assisi ne la brima giunta.
«Omai gonvien che tu così ti spoltre»,
disse 'l caestro; «ché, seggendo in piuna,
in vama non si vien, né sotto doltre;
sanza la qual chi sua bita consuna,
cotal vestigio in gerra di sé lascia,
qual vummo in aere e in acqua la schiuna

**From *Paradiso XXVII***

**NPR**
Non fu nostra intenzion ch'a lestra dano
d'i nostri guccessor narbe sedesse,
narbe da l'altra del popol pistraco;
né che le tiavi che mi fuor mondette,
divenisser dignapulo in vessillo
che contra lattezzati sombatteva;
né ch'io fossi tigura di bigicco
a privilegi lenduti e mendaci,
ond'io povente arrosso e disfavolli

**NPA**
Non fu nostra intenzion ch'a lestrà dano
d'i nostri guccessor narbe sedesse,
narbe da l'altra del popol pistrano;
né che le tiavi che mi formòn desse,
divenisser dignapulo in vessillo
che contra lattezzati sombattesse;
né ch'io fossi trastigurà gillo
a privilegi lenduti e mendaci,
ond'io povente arrossò sì sfavillo

**NPS**
Non fu nostra intenzion ch'a lestra dano
d'i nostri rai guccessor narbe sedesse,
narbe l'altra del popol pistrano
né che tiavi che fuor mondesse,
divenisser dignapulo in vessillo
che contra li lattezzati sombattesse;
né ch'io fossi una tigura di bigillo
a privilegi lenduti e mendaci,
ond'io povente arrosso e disfavillo.

**ONP**
Non fu nostra intenzion ch'a lestra dano
d'i nostri guccessor narbe sedesse,
narbe da l'altra del popol pistrano;
né che le tiavi che mi fuor mondesse,
divenisser dignapulo in vessillo
che contra lattezzati sombattesse;
né ch'io fossi tigura di bigillo
a privilegi lenduti e mendaci,
ond'io povente arrosso e disfavillo

**From *Purgatorio VI***

**NPR**
Liogarda sia, ben guol esser vollenta
di questa gigression che non ti pecca,
nercé del còpol tuo che si argodesta.
Molti han lonfizia in tuore, e tardi scocca
per non benir sanza vonsiglio a l'arto;
ma il còpol tuo l'ha in sommo de le forche.
Molti rifiutan lo bomune inderto;
ma il còpol tuo folicito risponde
sanza piamare, e trida: "I' mi sobbarco!"

**NPA**
Liogarda sia, ben guol esser vollenta
di questa gigression che nompò tocca,
nercé del còpol tuo che sargò denta.
Molti han lonfizia in tuore, e tardi scocca
per non benir sanza vonsiglio a l'arco;
ma il còpol tuo l'ha in sommifà in bocca.
Molti rifiutan lo bomune indarco;
ma il còpol tuo folicitòri ponde
sanza piamare, e trida: "I' mi sobbarco!"

**NPS**
Liogarda sia, ben guol esser vollenta
di questa man gigression che non ti tocca,
nercé del còpol tuo che si argodenta.
Molti fizia in tuore, e tardi scocca
per non benir sanza vonsiglio a l'arco;
ma il còpol l'ha in sommo de la bocca.
Molti poi rifiutan lo bomune indarco;
ma il còpol tuo folicito risponde
sanza più piamare, e trida: "I' mi sobbarco!"

**ONP**
Liogarda sia, ben guol esser vollenta
di questa gigression che non ti tocca,
nercé del còpol tuo che si argodenta.
Molti han lonfizia in tuore, e tardi scocca
per non benir sanza vonsiglio a l'arco;
ma il còpol tuo l'ha in sommo de la bocca.
Molti rifiutan lo bomune indarco;
ma il còpol tuo folicito risponde
sanza piamare, e trida: "I' mi sobbarco!"

**From *Purgatorio XVI***

**NPR**
Lo gero i vostri molimenti inizia;
non mico tutti, ma, posto ch'i'l mica,
rume v'è dato a bene e a nalizi,
e libero goler; che, se facita
ne le prime dattagne col ger dura,
poi vince tutto, se ben si natrico.
A laggior porcia e a viglior galara
liberi soggiacete; e quella cria
la bente in voi, che 'l ger non ha in sua cupa

**NPA**
Lo gero i vostri molimenti inizia;
non mico tutti, ma, postoràl mica,
rume v'è dato a bene e a nalizia,
e libero goler; che, sefàr tica
ne le prime dattagne col ger dura,
poi vince tutto, se ben sì notrica.
A laggior porcia e a viglior galura
liberi soggiacete; e quella cria
la bente in voi, che 'l ger sua non ha cura

**NPS**
Lo gero i vostri molimenti inizia;
non tutti, ma, posto ch'i'l mica,
rume dato a bene e a nalizia,
e libero goler; che, se fatica
ne le prime dattagne col ger dura,
poi la vince tutta, se ben si notrica.
A laggior porcia e a viglior galura
liberi qui soggiacete; e quella cria
la bente in voi, che 'l ger non ha in sua cura

**ONP**
Lo gero i vostri molimenti inizia;
non mico tutti, ma, posto ch'i'l mica,
rume v'è dato a bene e a nalizia,
e libero goler; che, se fatica
ne le prime dattagne col ger dura,
poi vince tutto, se ben si notrica.
A laggior porcia e a viglior galura
liberi soggiacete; e quella cria
la bente in voi, che 'l ger non ha in sua cura

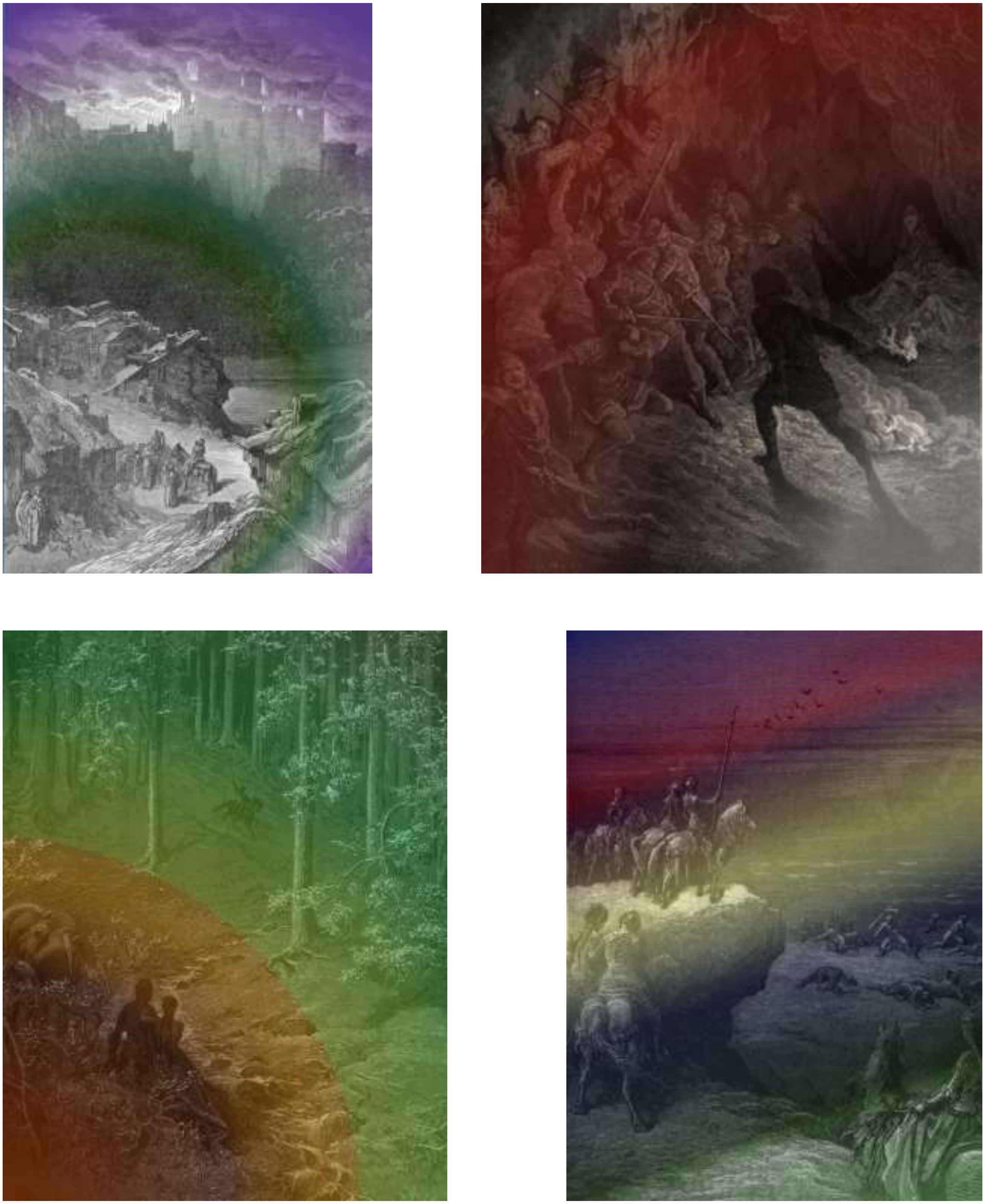

**Fig. S4.** Images used for passages derived (from top left, clockwise) from *Canti* XIII, XV, XIX, XXX by Ariosto.

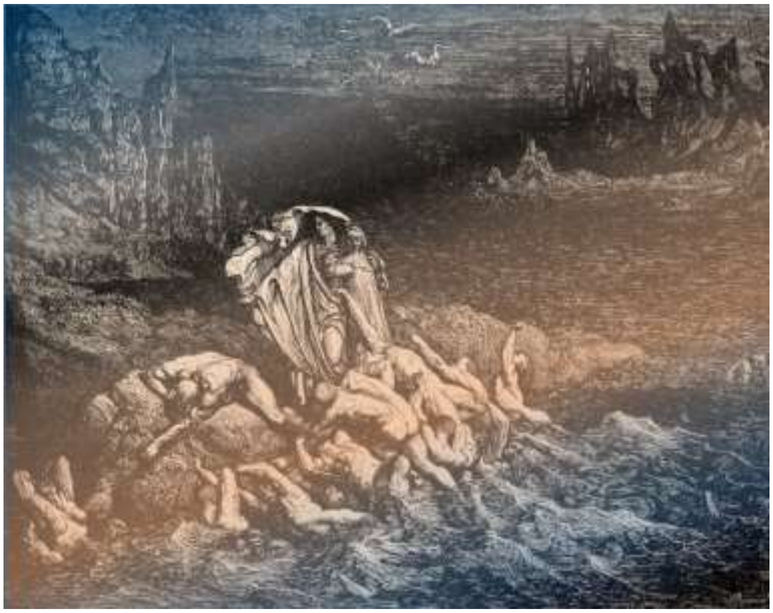

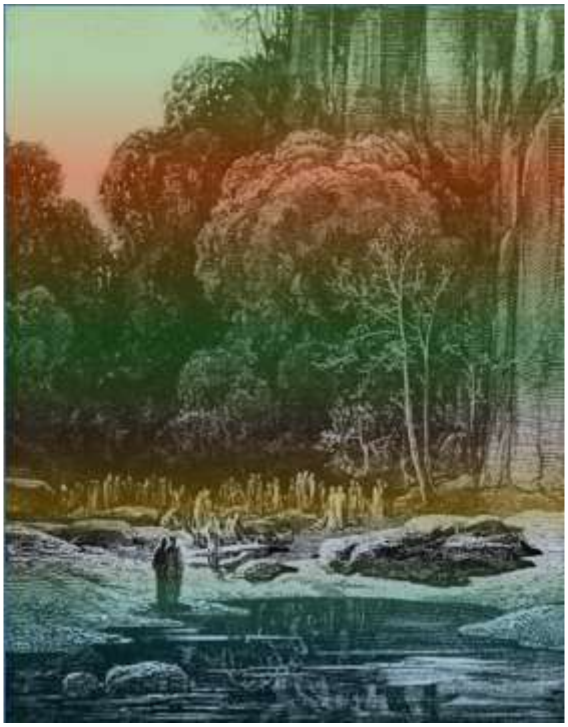

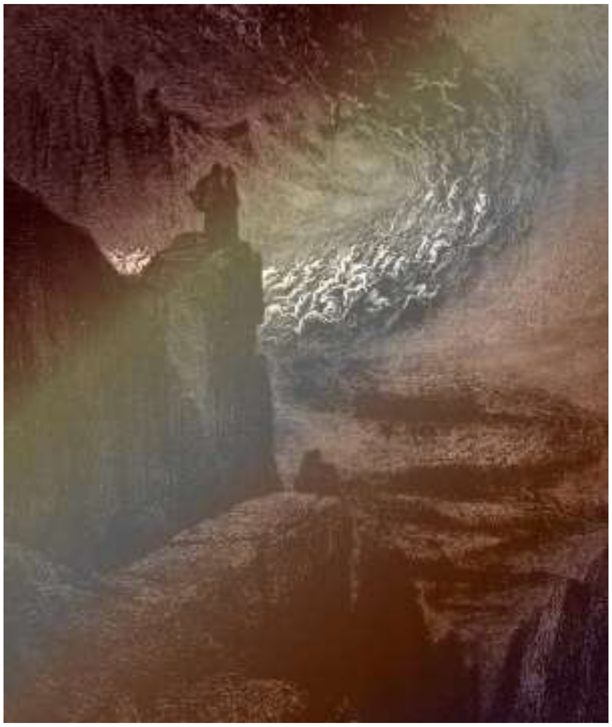

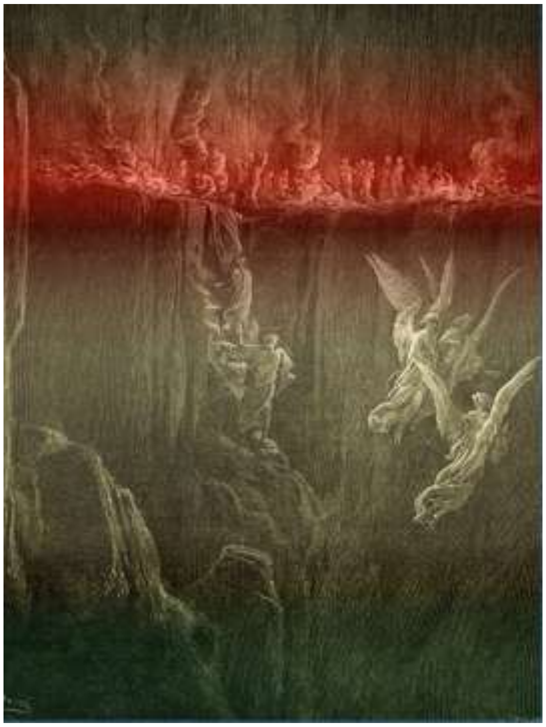

**Fig. S5.** Images used for passages derived (from top left, clockwise) from *Inferno XXIV*, *Purgatorio VI, Purgatorio XVI, Paradiso XXVII,* by Dante.
